# Drug-like Fragments Inhibit *agr-*Mediated Virulence Expression in *Staphylococcus* aureus

**DOI:** 10.1101/538173

**Authors:** Ian F. Bezar, Ameya A. Mashruwala, Jeffrey M. Boyd, Ann M. Stock

## Abstract

In response to the increasingly problematic emergence of antibiotic resistance, novel strategies for combating pathogenic bacteria are being investigated. Targeting the *agr* quorum sensing system, which regulates expression of virulence in *Staphylococcus aureus*, is one potentially useful approach for combating drug-resistant pathogens that has not yet been fully explored. A previously published study of a fragment screen resulted in the identification of five compound fragments that interact with the DNA-binding domain of the response regulator AgrA from *S. aureus*. We have analyzed the ability of these compounds to affect *agr*-mediated virulence gene expression in *S. aureus* cells. Three of the compounds demonstrated the ability to reduce *agr*-driven transcription of at the P2 and P3 promoters of the *agr* operon and increase biofilm formation, and two of these compounds also showed the ability to reduce levels of secreted toxins. The finding that the compounds tested were able to reduce *agr* activity suggests that they could be useful tools for probing the effects of *agr* inhibition.

Furthermore, the characteristics of compound fragments make them good starting materials for the development of compound libraries to iteratively improve the inhibitors.

## Introduction

*Staphylococcus aureus* is a dangerous human pathogen and a leading cause of endocarditis, bone and joint infections, pulmonary infections, and bacteremia^1^. *S. aureus* infections have become increasingly difficult to treat due to the growing prevalence of antibiotic-resistant strains. Methicillin-resistant *S. aureus* (MRSA) strains such as USA300 have become the predominant source of soft-tissue infections in the USA^2,3^. MRSA infections are often treated with vancomycin as a last-resort antibiotic; however, strains resistant to vancomycin have been reported^4,5^. Although clinical observation of vancomycin resistance in infections has been relatively limited, the threat highlights the urgent need for novel antibiotic therapies^6^.

In response to the problem of increasing antibiotic resistance, targeting bacterial virulence rather than viability has been proposed. Because virulence expression and regulation are important for the establishment and maintenance of an infection but are otherwise non-essential, it has been argued that targeting virulence might be less likely to lead to the development of resistance^7,8^. While these potential advantages make the idea of targeting virulence extremely appealing, this strategy remains largely untested.

In *S. aureus*, the *agr* quorum sensing system plays a major role in the regulation of virulence^9^. The *agr* system coordinates the timing of the transition to an invasive mode that entails increased production of virulence factors and a reduction in surface proteins^10^. Infection models have shown that disruption of the timing of *agr* activation leads to the attenuation of an infection^11^. The importance of *agr-*mediated expression of virulence genes has also been demonstrated in several infection models where *agr*-deficient strains generate significantly milder infections than their wild-type (WT) counterparts^12-15^.

The *agr* operon consists of four genes: *agrB*, *agrD*, *agrC*, and *agrA* that encode the components of the quorum sensing system^16^. Transcription of the operon is driven by the P2 promoter, which is activated by the response regulator AgrA in an autoregulated fashion. *agrD* encodes a 46-amino acid pro-peptide that is processed and secreted from the cell by the transmembrane endopeptidase AgrB^17,18^. The mature secreted AgrD is the auto-inducing peptide (AIP) of the quorum sensing system, which, after building up to sufficient extracellular concentrations, is capable of activating the receptor histidine kinase AgrC^19^. Activated AgrC promotes the transfer of a phosphoryl group to the response regulator AgrA, which in turn activates transcription at the P2 promoter, completing the auto-regulatory loop^20^. Phosphorylated AgrA also promotes transcription at the P3 promoter, leading to expression of RNAIII, a 514-nucleotide RNA molecule that both serves as the transcript for the *hld* gene encoding δ-hemolysin and functions as a regulatory RNA^21,22^. RNAIII plays a central role in effecting the transition to a virulent mode as it serves to enhance the expression of genes encoding toxins such as *hla* (α-hemolysin) while reducing the expression of genes encoding surface proteins, such as *spa* (protein A). The down-regulation of adhesion molecules upon the activation of the *agr* system is accompanied by the increased expression of enzymes capable of dissolving biofilm matrices, such as nucleases and proteases. Thus, increased *agr* activity results in the inhibition of biofilm formation as well as facilitating the dispersal of bacteria from pre-formed biofilms^23,24^.

AgrA is a response regulator of the LytTR family, characterized by a DNA-binding domain that is relatively uncommon among bacteria and absent from higher organisms^25^. LytTR domains are typically found in transcription factors that regulate virulence gene expression^26^. A previously conducted fragment screen against the AgrA LytTR domain identified five compounds that interacted with the DNA-binding domain at a common site that overlapped the DNA-binding surface. Three compounds were shown to inhibit interactions of the AgrA DNA-binding domain with its target DNA in an *in vitro* assay^27^. Drug-like fragments, which are smaller than typical small-molecule drugs and thus bind with relatively low affinity, are considered to be good starting points in drug-development pipelines^28^. We aimed to test the hypothesis that the previously identified fragments, which target a DNA-interaction surface of AgrA, would inhibit AgrA activity in *S. aureus* cells. Here we present data demonstrating that several of the compounds identified in the original screen inhibit virulence gene activation in *S. aureus* in ways that are consistent with inhibition of the *agr* system. These data suggest that these molecules are not only useful for the study of effects of *agr* inhibition but also have potential as starting molecules for the design of improved inhibitors.

## Results

### Treatment with inhibitors results in decreased activation of the P3 promoter

Phosphorylated AgrA drives transcription at the P3 promoter, leading to expression of RNAIII^22^. To test the ability of the inhibitory compounds to disrupt the activation of transcription by AgrA, we employed a cell-based reporter assay using a plasmid containing the *gfp* gene under the transcriptional control of the P3 promoter^29^.

Untreated WT cultures grown for 8 h demonstrated robust expression of GFP, indicating strong transcription of RNAIII (Fig. 1a). In contrast, cultures grown in the presence of compounds **1-4** at a concentration of 120 μM resulted in reduced expression of GFP (Fig. 1a). This concentration of the compounds did not inhibit growth (Supplementary Fig. S1). Treatment with compound **5** did not significantly alter reporter activity. The largest reduction in GFP expression was achieved with compound **1**, although levels were not reduced to those of the ∆*agr::tet* strain, henceforth referred to as the ∆*agr* strain. None of the compounds had significant effects on the ∆*agr* strain (p>0.05), suggesting that the compounds exert their inhibitory effects via the *agr* system.

**Figure 1.**
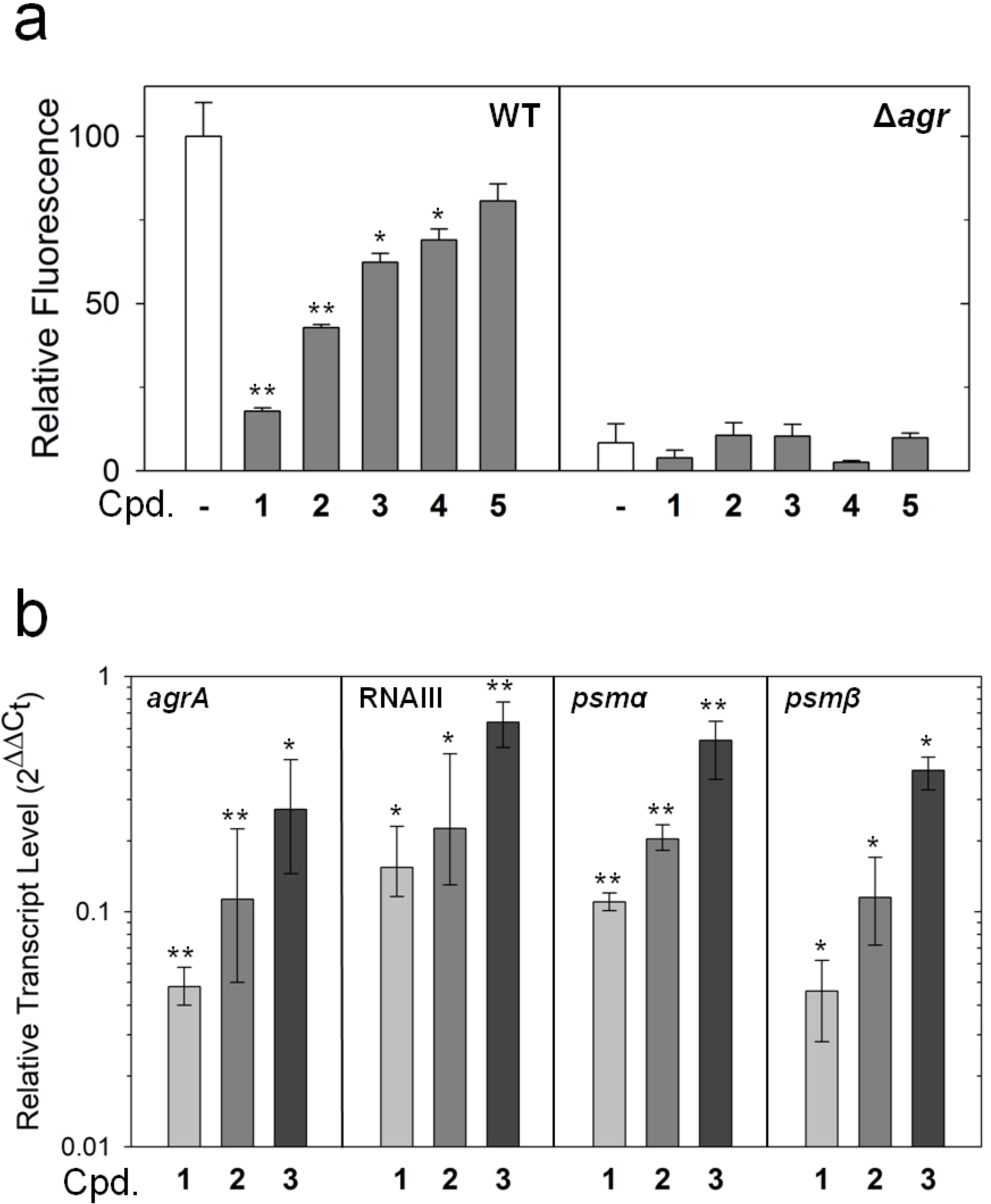
AgrA-driven transcription is inhibited by compounds. (**a**) Plasmid-based GFP expression driven by the P3 promoter is inhibited by compounds. Cultures were grown for 8 h in the presence of the indicated compound (Cpd.) at a concentration of 120 μM or DMSO alone (-). Bars represent OD-normalized fluorescence averaged from three separate experiments relative to the untreated sample (normalized to 100) with error bars indicating standard deviations. Statistical significance relative to the sample without compound was determined using a Student’s t-test (* p<0.05, ** p<0.01). (**b**) Compounds reduce transcript levels of genes directly regulated by AgrA. mRNA was isolated for qPCR analysis after cultures were grown for 8 h in the presence of the indicated compound (Cpd.) at a concentration of 120 μM or DMSO alone. Bars represent the fold change in mRNA level from treated cultures relative to untreated cultures averaged from three separate experiments. Statistical significance and 95% confidence intervals (displayed as error bars) were determined using REST2009 software^59^ (* p<0.05, ** p<0.01).

Compounds **1-3**, which showed the greatest inhibitory effects, were chosen for further analyses. Treatment with compounds **1-3** had no effects on GFP expression in strains containing transcriptional reporter plasmids for *dps* and *recA*, two genes that are not known to be under *agr* regulation^30^ (Supplementary Fig. S2). These data suggest that the compounds do not act generally to reduce GFP expression and are consistent with the interpretation that the compounds reduce transcription from the P3 promoter through the *agr* system.

### Treatment with inhibitors reduces levels of transcripts directly regulated by AgrA

AgrA directly regulates the transcription of several genes involved in virulence regulation. It promotes the transcription of genes encoding the phenol-soluble modulin (PSM) α and β proteins^30^, the *agrBDCA* operon at the P2 promoter, and the regulatory RNA, RNAIII, at the P3 promoter. Inhibition of AgrA is therefore expected to result in reduced cellular levels of these AgrA-regulated transcripts. Quantitative real-time PCR was performed to determine the transcript levels of *psmα1*, *psmβ1*, *agrA*, and RNAIII in cultures grown with each compound at a concentration of 120 µM for 8 h. Levels of each of the tested transcripts were significantly lower in cultures treated with compounds **1-3** compared to those in untreated cultures (between 1.6-22 fold decrease in expression) (Fig. 1b). The reduction of transcript levels upon treatment with the compounds is consistent with the interpretation that the compounds interfere with regulation of specific transcripts by AgrA.

### Treatment with inhibitors results in reduced cellular levels of AgrA

AgrA drives transcription of the *agrBDCA* operon at the P2 promoter. Inhibition of AgrA would therefore be expected to result in reduced cellular levels of AgrA. In order to test whether the compounds reduced AgrA levels, western blots were employed to measure levels of AgrA in a ∆*spa* strain. The ∆*spa* strain was used to avoid interference from the immunoglobulin-binding protein A with western blotting. Compounds were added to cultures upon reaching an OD of ~2.5, designated as time zero, at which AgrA levels were moderate but detectable. Samples were collected after an additional 8 h of growth, a point expected to exhibit high levels of AgrA, induced by high cell density (Supplementary Fig. S3). In untreated samples, levels of AgrA approximately doubled in the 8-h growth window (Fig. 2a). However, treatment with compounds **1-3** resulted in a concentration-dependent reduction in levels of AgrA. Importantly, levels of AgrA from cultures treated with each of the three compounds at concentrations of 250 μM did not increase significantly during the 8-h growth period, indicating nearly complete inhibition of new AgrA synthesis.

**Figure 2.**
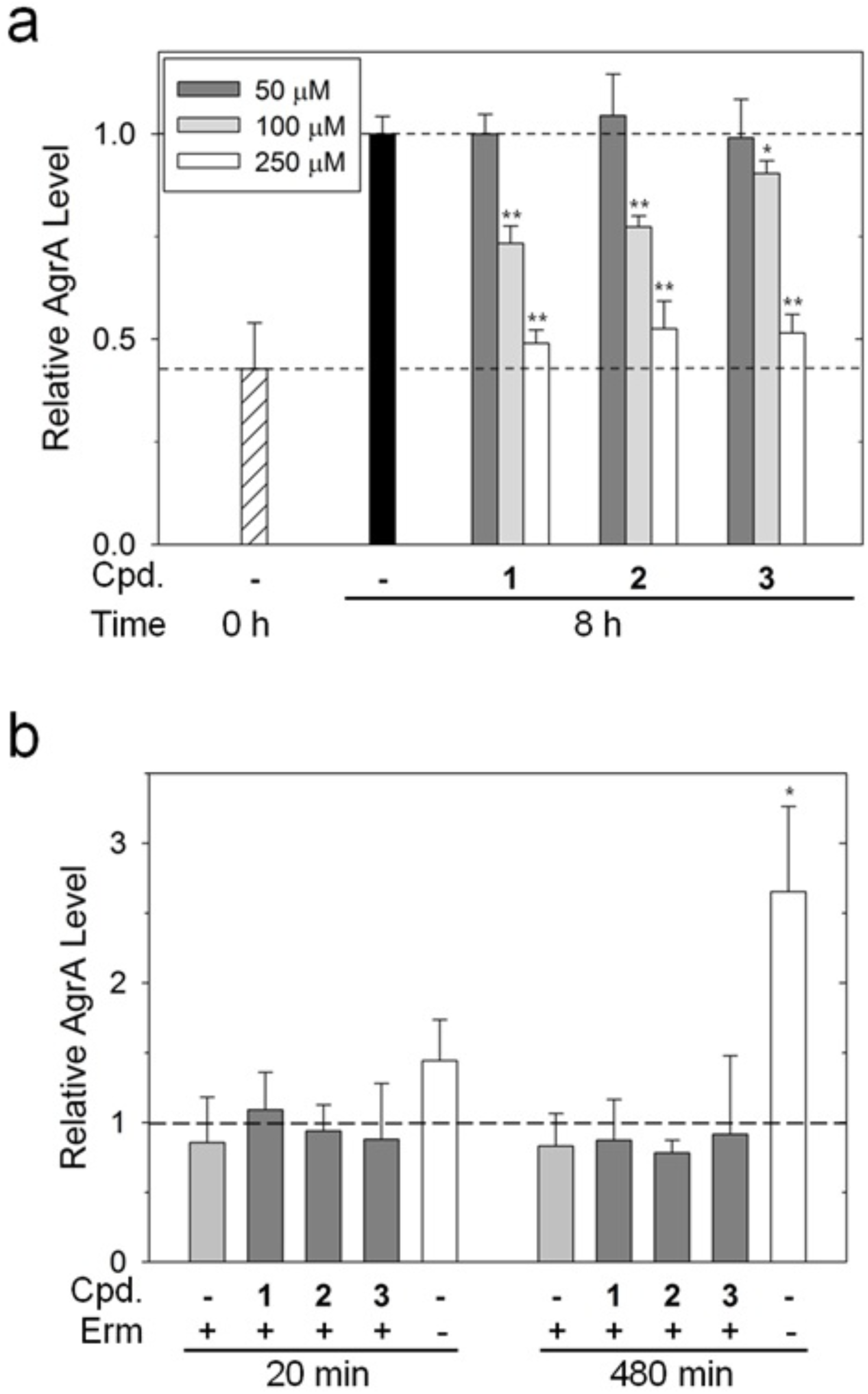
Compounds reduce levels of AgrA without altering AgrA stability. (**a**) Cultures of USA300 *spa::kan* were grown to OD 2.5 (represented by t=0 and the lower dashed line) at which time the cultures were divided and compounds were added at the indicated concentrations. Samples were harvested after 8 h of growth and AgrA levels were determined by western blotting. The results of three separate blots (one for each compound) analyzed in parallel are shown. Bars indicate densitometry quantitation of AgrA band intensities averaged from three independent experiments normalized to the 8-h sample without compound (-, DMSO only, represented by the upper dashed line) with error bars representing standard deviations. (**b**) Cultures of USA300 *spa::kan* were grown to OD 2.0, at which time samples were taken and erythromycin was added to a concentration of 10 μM. Additional samples were taken upon addition of compounds at 20 min of incubation with erythromycin and at 480 min of incubation. The results of three separate blots (one for each compound) analyzed in parallel are shown. The level of AgrA before the addition of erythromycin is depicted by the dashed line. Bars represent the averages of three replicates normalized to untreated cultures (t=0) for samples collected at 20 and 480 min with standard deviations shown as error bars. Statistical significance relative to the samples of erythromycin-treated cultures without compound (t=20, 480) was determined using a Student’s t-test (* p<0.05, ** p<0.01).

An alternate explanation for the reduction in AgrA levels is that treatment with the compounds led to an increase in the turnover of AgrA. To examine the effects of the compounds on protein stability, cultures were pre-treated with the translational inhibitor erythromycin prior to treatment with compounds **1-3**. Cultures were sampled immediately prior to the addition of erythromycin, 20 min post addition of the compounds, and finally after 8 h of incubation. In the absence of erythromycin, levels of AgrA increased over time, while cultures grown with erythromycin maintained constant levels of AgrA (Fig. 2b). The addition of compounds **1-3** to samples pre-treated with erythromycin resulted in no significant difference in the levels of AgrA. These results are consistent with the interpretation that the compounds disrupt *agr* activity by interfering with the ability of AgrA to activate *agrBDCA* transcription.

### Treatment with inhibitors results in decreased production of exoproteins, reduced hemolytic activity, and altered levels of the *spa* and *hla* transcripts

The *agr* system regulates virulence in part by increasing the expression of exoproteins such as hemolytic toxins^31^. Inhibition of AgrA activity would therefore be expected to decrease exoprotein production and lead to decreased hemolytic activity. The presence of exoproteins was analyzed in the spent media supernatant of cultures grown over a period of 8 h. Exoprotein levels of the culture supernatant obtained from WT cultures treated with compounds 1 and 2 were visibly decreased relative to those from untreated cultures when analyzed using SDS-PAGE (Fig. 3a). In addition, differences in the relative intensities of some bands were observed. No appreciable change in exoprotein profiles was noted in the *∆agr* strain upon treatment with the compounds.

**Figure 3.**
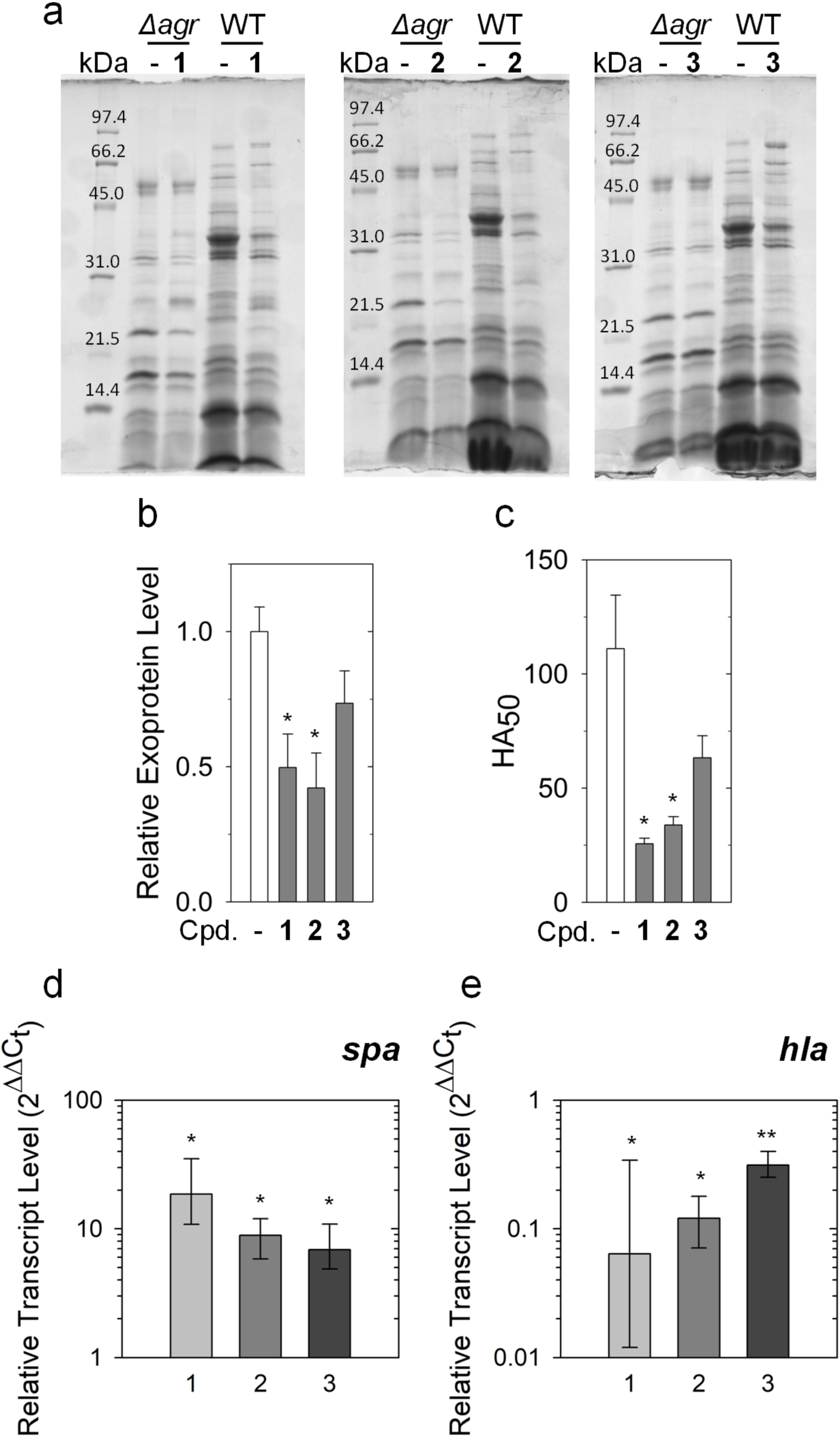
Expression of exoproteins and surface proteins in *S. aureus* is altered by compounds. Cultures of WT and ∆*agr* were grown in the absence or presence of compounds at a concentration of 120 μM for 8 h of post-treatment growth. (**a**) Secreted proteins were isolated from culture media and samples were analyzed using SDS-PAGE. (**b**) The concentration of total secreted protein was determined using a Bradford protein assay. Levels of *agr*-dependent protein secreted by WT in the absence and presence of compounds were estimated by subtracting the level of secreted protein in ∆*agr* from the average levels in WT determined from three replicates. Bars represent values relative to the untreated sample (normalized to 1) with errors bars representing standard deviations. (**c**) Hemolytic activity was assessed after 8 h of growth by adding dilutions of filtered culture media to defibrinated rabbit blood and measuring liberated hemoglobin. HA_50_ values were calculated and are shown with error bars representing the standard error of the mean. (**d-e**) Compounds alter transcript levels for *spa* (**e**) and *hla* (**f**). mRNA was isolated for qPCR analysis after cultures were grown for 8 h in the presence of the indicated compound at a concentration of 120 μM or DMSO alone. Bars represent the fold change in mRNA level from treated cultures relative to untreated cultures averaged from three separate experiments. Statistical significance and 95% confidence intervals (displayed as error bars) were determined using REST2009 software (* p<0.05, ** p<0.01).

To quantify the differences in levels of secreted proteins upon treatment with compounds, total protein levels in the culture media were measured using a Bradford protein assay. Secreted proteins were significantly reduced (p<0.05) in WT cultures treated with compound **1** and compound **2**, but not compound **3** (Fig. 3b). No appreciable change in protein levels was observed when ∆*agr* cultures were treated with compounds (Supplementary Fig. S5). Similarly, treatment of the cultures with compounds **1** and **2**, but not **3**, resulted in a significant decrease in the hemolytic activity of culture supernatants (p>0.05) (Fig. 3c).

While expression of RNAIII alters expression of protein A and α-hemolysin via modulation of translation^22^, down-stream effects of activation of *agr* system also lead to altered transcription of the *spa* and *hla* genes^30,32^. The effects of the compounds on the transcription of the *spa* and *hla* genes were assessed using quantitative real-time PCR. Addition of compounds **1-3** to cultures resulted in increased levels of *spa* transcripts (Fig. 3d) and reduced levels of *hla* transcripts (Fig. 3e).

### Treatment with inhibitors promotes biofilm formation

Activation of the *agr* system inhibits the formation of biofilms and promotes the dispersal of biofilm matrices. Consequently, strains with inactive *agr* systems demonstrate increased formation of biofilms^23^. Therefore, inhibition of *agr* activity was expected to promote biofilm formation. To assess levels of biofilm formation, cultures were incubated in 96-well plates without shaking. While untreated cultures developed moderate biofilms, treatment with compounds **1-3** resulted in increased biofilm formation in a concentration-dependent manner (Fig. 4a-c). Interestingly, treatment with compound **1** at a concentration of 125 µM resulted in biofilm formation at a level comparable to that of the ∆*agr* strain, suggesting that this compound is capable of completely reversing AgrA-mediated repression of biofilm formation (Fig. 4d).

**Figure 4.**
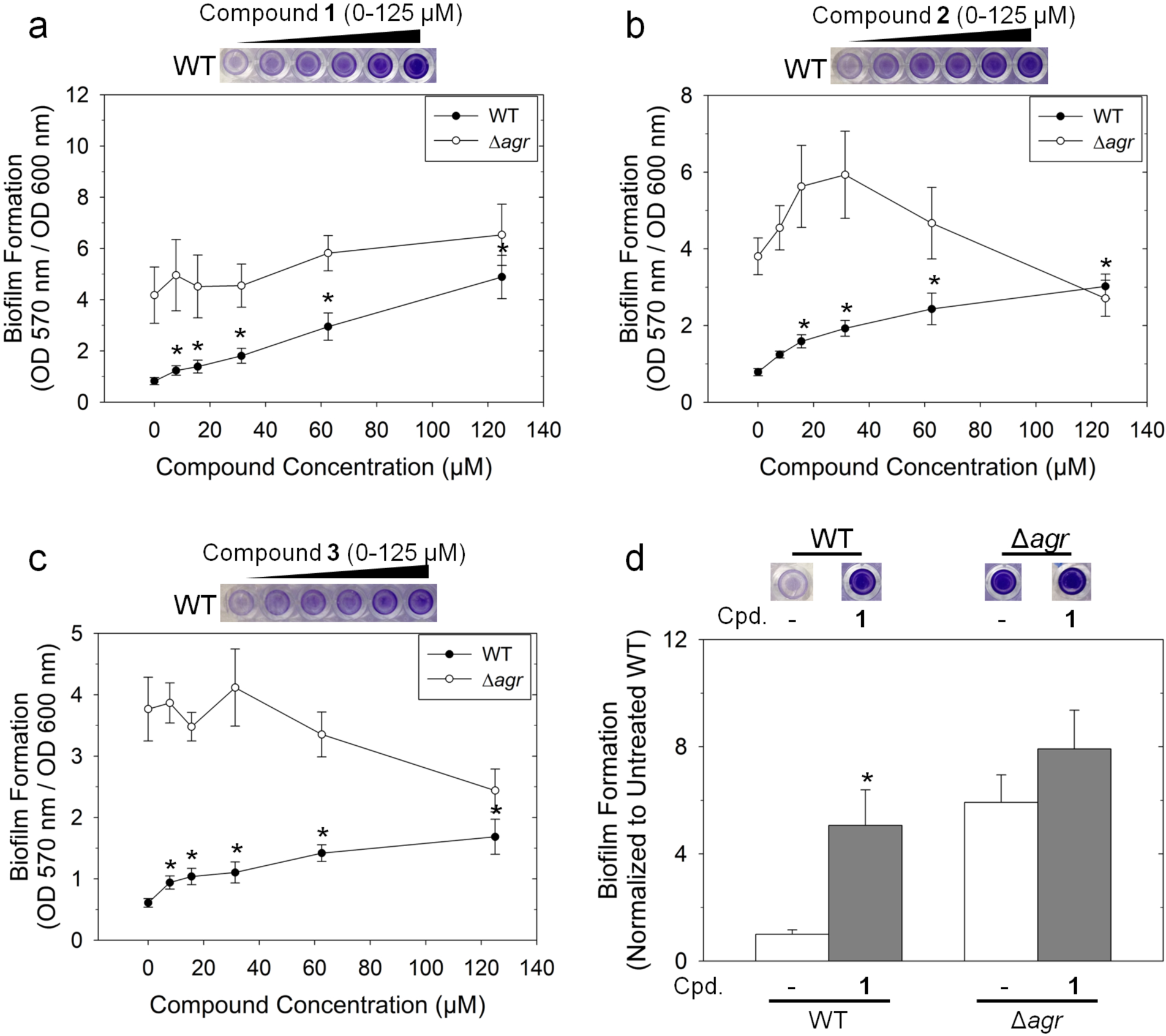
Treatment with compounds promotes biofilm formation. (**a-c**) WT and ∆*agr* cultures were grown statically in the presence of compounds to promote biofilm formation. After staining plates with crystal violet, biofilms were quantified by measuring absorbance at 570 nm. Bars and error bars represent absorbances and standard deviations of 6 replicates, respectively. (**d**) Biofilm formation of WT and ∆*agr* cultures treated with compound **1** at a concentration of 120 µM are shown normalized to the untreated WT sample with error bars representing standard deviations from 6 replicates. Statistical significance relative to the untreated sample for each strain was determined using a Student’s t-test (* p<0.05). Representative images of stained wells are included in each panel.

### Treatment with inhibitors results in decreased production of exoproteins in strains of different *agr* types

Across different strains of *S. aureus*, a region of hyper-variability exists within the *agr* operon, encompassing the latter half of *agrB, agrD* and the first half of *agrC*^33^. The resultant differences in AgrB, AgrD, and the *N*-terminal domain of AgrC allow for the production (in the case of AgrB and AgrD) and recognition (in the case of AgrC) of different autoinducing peptides. Based on these differences, *S. aureus* strains are classified as one of four *agr* types, numbered I-IV^34,35^. In contrast to *agrB*-*C*, the sequence of *agrA* is highly conserved within *S. aureus*, and therefore inhibition of AgrA would be expected to affect strains of all *agr* types. The USA300 LAC strain belongs to group I. We examined exoprotein levels in type II, III, and IV strains. Treatment with compounds **1** and **2** resulted in decreased production of exoproteins in N315 (*agr* type II), MW2 (*agr* type III), and MN EV (*agr* type IV) strains (Fig. 5). These data are consistent with the proposed mechanism whereby the inhibitors are functioning primarily through inhibition of AgrA.

**Figure 5.**
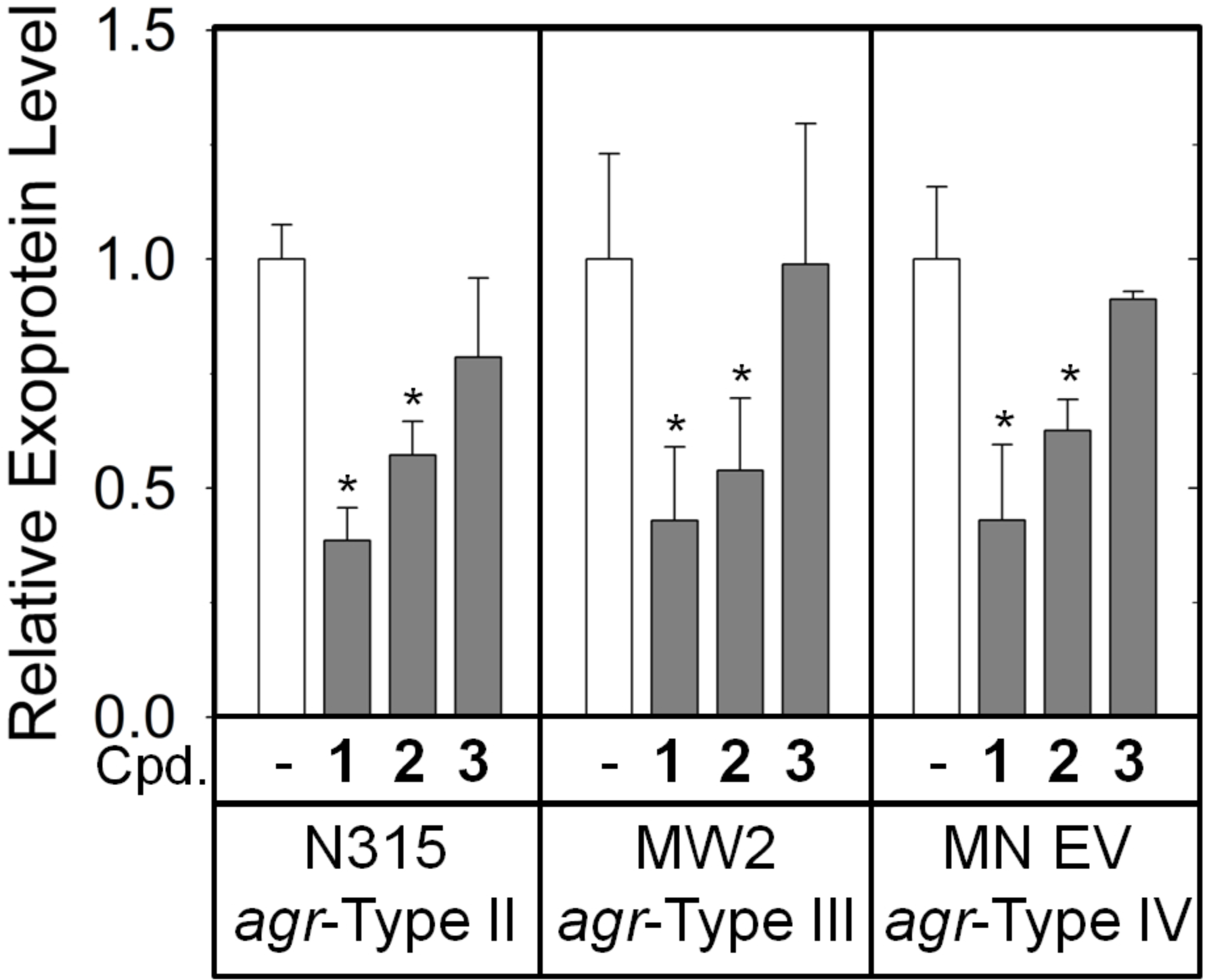
Treatment with compounds reduces exoprotein expression among *S. aureus* strains of *agr*-types II-IV. The concentration of total protein secreted into culture media was determined using a Bradford assay, and levels were normalized to those of untreated samples. Statistical significance relative to the untreated samples was determined using a Student’s t-test (* p<0.05).

### Compounds interact with AgrA with sub-millimolar affinity

Binding of the compounds to AgrA_C_ was previously demonstrated by NMR WATERGATE W5 LOGSY and chemical shift perturbation analyses^27^, however, binding affinities were not directly determined. Isothermal titration calorimetry (ITC) was used to measure the affinity of compounds **1-3** for AgrA_C_. Isotherms were easily fitted to one-binding-site models. Fitting of the isotherms was improved by fixing the stoichiometry to N=1. Fitting of the isotherm generated by compound 1 resulted in a measured ∆H of - 1154 ± 487.0 cal mol^−1^, ∆S of 23.4 cal mol^−1^deg^−1^, and a *K*_d_ of 485 ± 39.3 μM (Fig. 6a). Fitting of the isotherm generated by compound **2** yielded a measured ∆H of −423.4 ± 25.19 cal mol^−1^, ∆S of 14.0 cal mol^−1^deg^−1^, and a *K*_d_ of 417 ± 62.9 μM (Fig. 6b). Fitting of the isotherm generated by compound **3** yielded a ∆H of −340.0 ± 35.53 cal mol^−1^, ∆S of 17.0 cal mol^−1^deg^−1^, and a *K*_d_ of 110 ± 45.6 μM (Fig. 6c). The relatively large errors associated with the fitting of these parameters can be attributed to difficulties in analyzing low-affinity interactions.

**Figure 6.**
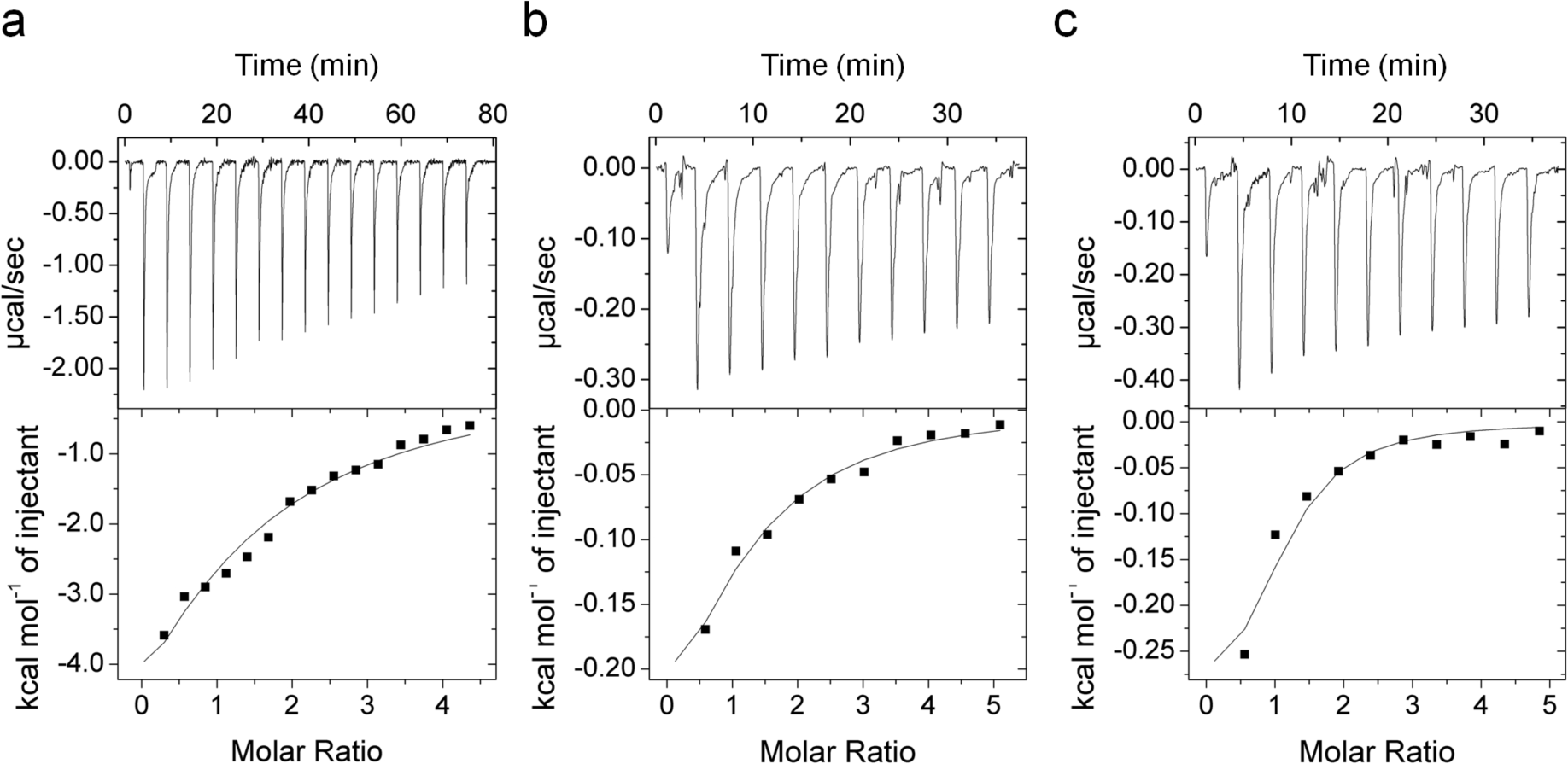
Compounds bind to AgrA_C_ with sub-millimolar affinity. ITC experiments were performed by titrating (**a**) compound **1** (10.0 mM) into 375 μM AgrA_C_, (**b**) compound **2** (10.0 mM) into 375 μM AgrA_C_, and (**c**) compound **3** (5.0 mM) into 250 μM AgrA_C_. After subtraction of the heat of dilution, isotherms were fitted to a one-binding-site model to generate thermodynamic parameters.

## Discussion

Inhibition of virulence has been proposed as a strategy for combating bacterial infections because it exploits previously unexplored targets for inhibition and also has the potential to limit selection for resistance. *S. aureus* is one pathogen for which virulence inhibition is especially appealing as it produces an extensive array of harmful virulence factors^36^ and has a history of antibiotic resistance that dates back to the initial introduction of penicillin^37^. Within *S. aureus*, the *agr* quorum sensing system has been specifically identified as a potential target for therapeutic development because of the central role the system plays in regulating virulence gene expression. The potential of this strategy is supported by studies using certain infection models that demonstrate attenuated infections with *agr*-deficient strains^11,13,14^. Despite the recent interest in the strategy of targeting virulence, both the effectiveness of using a therapeutic agent to target virulence and the reduction in selective pressure caused by such agents remain unproven.

Because of the unusual three-dimensional fold of the AgrA DNA-binding domain and the key role that AgrA plays in both virulence activation (by promoting expression of toxins) and regulation (by activating the quorum sensing mechanism), AgrA has become a target of interest for inhibiting virulence^38,39^. Our previous fragment screen against the AgrA DNA-binding domain identified five compounds with the potential to act as inhibitors of AgrA^27^. While these compounds were conclusively shown to interact with the LytTR domain of AgrA, they were unproven in their abilities to inhibit *agr* activity in *S. aureus* cells.

AgrA functions as a transcription factor, and thus the immediate effect of inhibiting AgrA would be to alter the levels of AgrA-regulated transcripts. The presented studies demonstrate that compounds **1-4** significantly reduced the expression of GFP driven by the P3 promoter, and compounds **1-3** reduced levels of transcripts of *agrA*, RNAIII, *psmα*, and *psmβ,* strongly suggesting that the compounds are acting to interfere with AgrA-regulated transcription. However, while the reduction in transcript levels were significant, they were not comparable to levels seen in the *∆agr* strain, which were below the threshold of detection under the tested conditions. Optimization of the compounds will therefore be required to achieve complete inhibition of AgrA.

The expression of RNAIII in particular is of central importance for the transition to virulence activation^40,41^. RNAIII coordinates virulence activation via interaction with multiple mRNA targets, resulting in both the direct regulation of some mRNAs and indirect regulation via modulation of expression of regulators such as Rot and MgrA^32,42-44^. Because RNAIII expression impacts several layers of virulence regulation, it substantially alters the expression profile of *S. aureus*, with studies suggesting that approximately 70 extracellular proteins are regulated by RNAIII^45^. Therefore, for an AgrA inhibitor to function as an inhibitor of virulence activation, expression of RNAIII-regulated virulence factors must be reduced. Consistent with decreased production of RNAIII, treatment with compounds **1-3** reduced levels of secreted proteins and *hla* transcript levels, and treatment with compounds **1** and **2** reduced hemolytic activity. Furthermore, treatment with compounds **1-3** promoted biofilm formation and increased levels of *spa* transcript. These effects are all consistent with a reduction in RNAIII production. However, the reductions in these downstream effects were not directly correlated with the reduction in RNAIII expression, likely reflecting the responsiveness of individual RNAIII-regulated systems to the level of RNAIII. In addition, compounds **1-3** were capable of significantly reducing the expression of AgrA itself, supporting the interpretation that the compounds were acting by inhibiting transcription at both the P2 and P3 promoters. By reducing both the autoregulatory and virulence factor production functions of AgrA, the potency of virulence inhibition could potentially be enhanced. Together, these results suggest that the compounds inhibit virulence gene activation in *S. aureus* cells in a manner that is consistent with inhibition of the *agr* system.

The compounds examined in this study inhibited several different aspects of *agr*-mediated virulence factor activation, suggesting that they could serve as useful tools for experimental validation of the strategy of targeting the *agr* system to combat *S. aureus* infections. Pressing questions regarding the strategy of targeting the *agr* system to combat *S. aureus* infections remain unanswered. One of the most problematic issues is the promotion of biofilm formation that is associated with a reduction in *agr* activity. Formation of biofilms is associated with an increase in antibiotic resistance^46^, and *agr* deficiency has been associated with types of infections for which biofilm formation is especially prevalent^47^. It is possible that the therapeutic efficacy of *agr* inhibition would depend on the type of infection being treated. Furthermore, the nature of the *agr* quorum sensing system suggests that the activation of virulence gene expression must be precisely timed in order to be effective. In fact, specifically timed transient inhibition has been shown to attenuate model infections^11^. In light of these timing requirements, it is likely that inhibition at a point either too late or too early during an infection may reduce or eliminate the efficacy of *agr* inhibition. Discovering the ideal timing of virulence factor inhibition is important for analyzing the effectiveness of the strategy. Finally, the question of whether or not inhibiting virulence results in selective pressure that leads to the development of resistance as seen with traditional antibiotics is still unknown. These questions may be more easily answered using chemical inhibitors that allow variation in both timing and dose, rather than by use of genetic mutations. Compound **1**, the strongest inhibitor we tested, reduced *agr* activity in many assays to levels close to those of an *agr* mutant, indicating that compound **1** may be almost as effective as using mutant strains while retaining the flexibility of using chemical inhibition.

Across all of the assays that were employed, a consistent pattern emerged where compound **1** was the most potent inhibitor followed by compounds **2** and **3**. It is interesting that that the affinity of compound **1** determined by ITC analyses was significantly lower than that of compounds **2** or **3**. The discrepancy between the strength of binding (where compounds **2** and **3** bound tighter than compound **1**) and the results from cellular assays (where compound **1** was consistently a stronger inhibitor than both compounds **2** and **3**) is likely explained by other characteristics of compound **1** that allow it to function better as an inhibitor either during uptake or within the cellular environment. Compounds **1-3 all** bind with *K*_d_’s in the range of 10^−4^ M. These modest binding affinities are typical of small compounds identified in fragment screens, with the expectation that affinities can be greatly increased as the compounds are built out^28^.

Recently, several attempts to develop inhibitors of the *agr* system have been pursued using different strategies. Approaches using AIP analogs^48^, identifying natural product inhibitors^49^, and using traditional chemical inhibition^50-55^ have yielded promising results. In particular, two compounds, Savirin^54^ and ω-Hydroxyemodin (OHM)^55^, are suspected of inhibiting the LytTR domain of AgrA and are likely to behave similarly to the inhibitory compounds reported in the present study. Both Savirin and OHM were shown to reduce *agr* activity within *S. aureus* and were also effective at reducing the severity of infections in mouse models^54,55^. Interestingly, OHM shares structural features with compound **1**: both the xanthene base of compound **1** and the anthraquinone OHM feature a similar three-ringed foundation. However, without experimental data to determine how each compound interacts with the AgrA LytTR domain, the importance of the structural similarities of the compounds to their inhibitory functions cannot be assessed.

All five of the compounds tested in these studies originated from a fragment screen library. Fragments are designed to explore large areas of chemical space and to serve as good starting points for the development of therapeutics as they are small in size and amenable to further chemical modifications. The three compounds that we have shown to substantially, albeit incompletely, inhibit *agr* activity in *S. aureus* cells are logical starting points for further development. However, it should be noted that our previous studies indicate that all five of the compounds bind to AgrA at a similar site, targeting a region of the LytTR domain that overlaps with a surface involved in DNA binding. Thus all compounds might be considered as potential candidates for future pursuit. Just as the affinities of compound **1** and compound **3** are not correlated with the strength of inhibition in cells, the differences observed in the potency of inhibition across all compounds could be due to many different characteristics that are likely to be altered as the size and the complexity of the fragments are increased to enhance affinity, specificity, and intracellular inhibitory activity.

## Materials and Methods

### Compounds used

9H-xanthene-9-carboxylic acid, 2-(4-methylphenyl)-1,3-thiazole-4-carboxylic acid, 4-hydroxy-2,6-dimethylbenzonitrile, 4-phenoxyphenol, and [5-(2-thienyl)-3-isoxazolyl]methanol were obtained from Sigma-Aldrich, MO (Table 1). Unless otherwise indicated, stock solutions were freshly prepared with 2.4 mM compound in 100% anhydrous dimethylsulfoxide (DMSO).

**Table 1:**
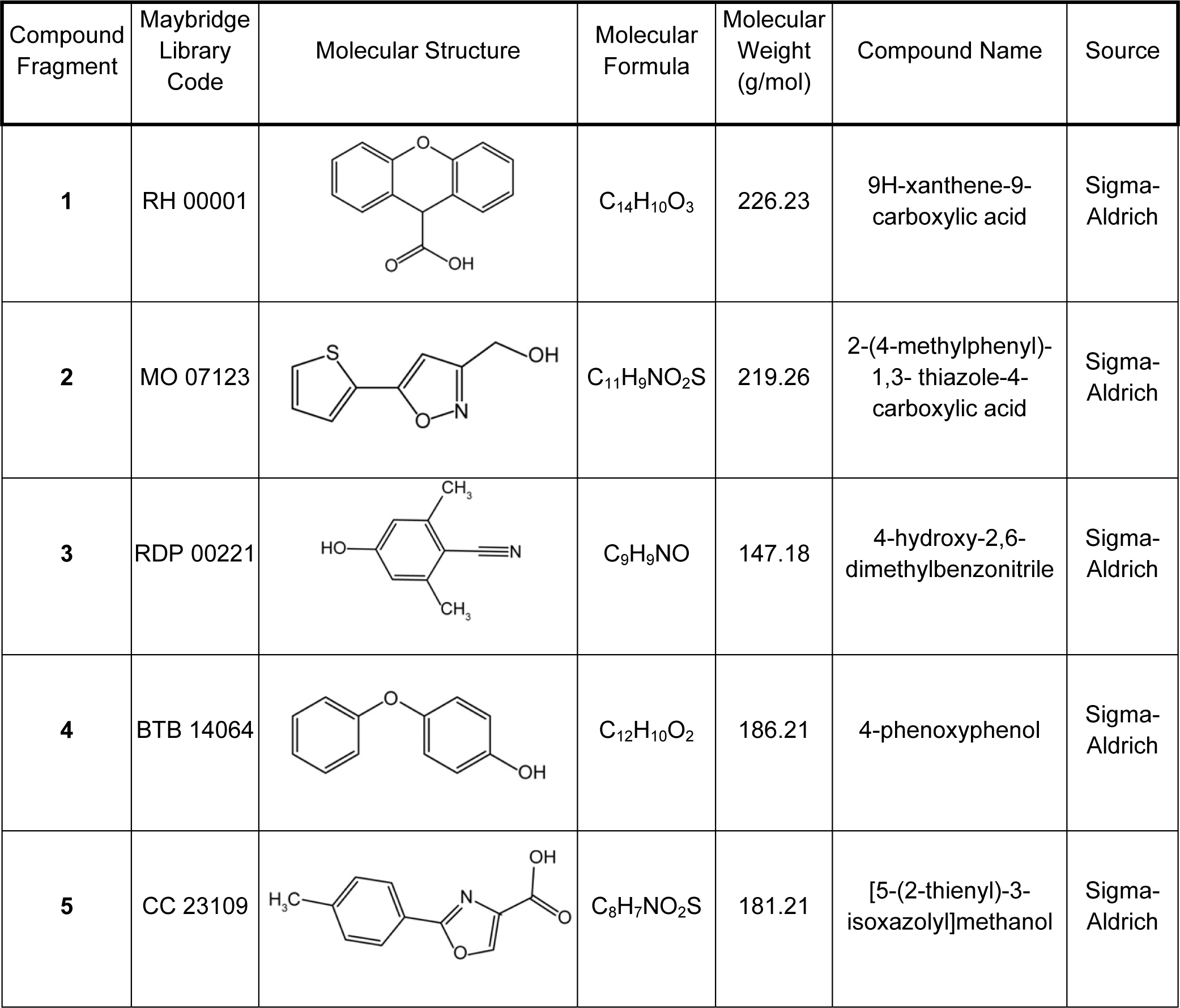
Compound Fragments

| Compound Fragment | Maybridge Library Code | Molecular Structure | Molecular Formula | Molecular Weight (g/mol) | Compound Name | Source |
| --- | --- | --- | --- | --- | --- | --- |
| 1 | RH 00001 |  | C<sub>14</sub>H<sub>10</sub>O<sub>3</sub> | 226.23 | 9H-xanthene-9-carboxylic acid | Sigma-Aldrich |
| 2 | MO 07123 |  | C<sub>11</sub>H<sub>9</sub>NO<sub>2</sub>S | 219.26 | 2-(4-methylphenyl)-1,3-thiazole-4-carboxylic acid | Sigma-Aldrich |
| 3 | RDP 00221 |  | C<sub>9</sub>H<sub>9</sub>NO | 147.18 | 4-hydroxy-2,6-dimethylbenzonitrile | Sigma-Aldrich |
| 4 | BTB 14064 |  | C<sub>12</sub>H<sub>10</sub>O<sub>2</sub> | 186.21 | 4-phenoxyphenol | Sigma-Aldrich |
| 5 | CC 23109 |  | C<sub>8</sub>H<sub>7</sub>NO<sub>2</sub>S | 181.21 | [5-(2-thienyl)-3-isoxazolyl]methanol | Sigma-Aldrich |

### Bacterial growth conditions

Unless otherwise stated, the *S. aureus* strains used in this study (Table 2) were constructed in the *S. aureus* community-associated USA300 strain LAC that was cured of the native plasmid pUSA03, which confers erythromycin resistance^56^. Unless specifically mentioned, *S. aureus* cells were cultured either using aerobic growth with a flask/tube headspace to culture medium volume ratio of 10:1 or in 96-well plates containing 200 μL total volume (detailed procedure below). Liquid cultures were grown at 37°C in Trypticase Soy Broth (TSB) with shaking at 200 rpm unless otherwise indicated. Difco Bacto agar was added (15 g L^−1^) for solid medium. For routine plasmid maintenance, liquid media were supplemented with chloramphenicol (10 μg mL^−1^) or erythromycin (3.3 μg mL^−1^).

**Table 2:**
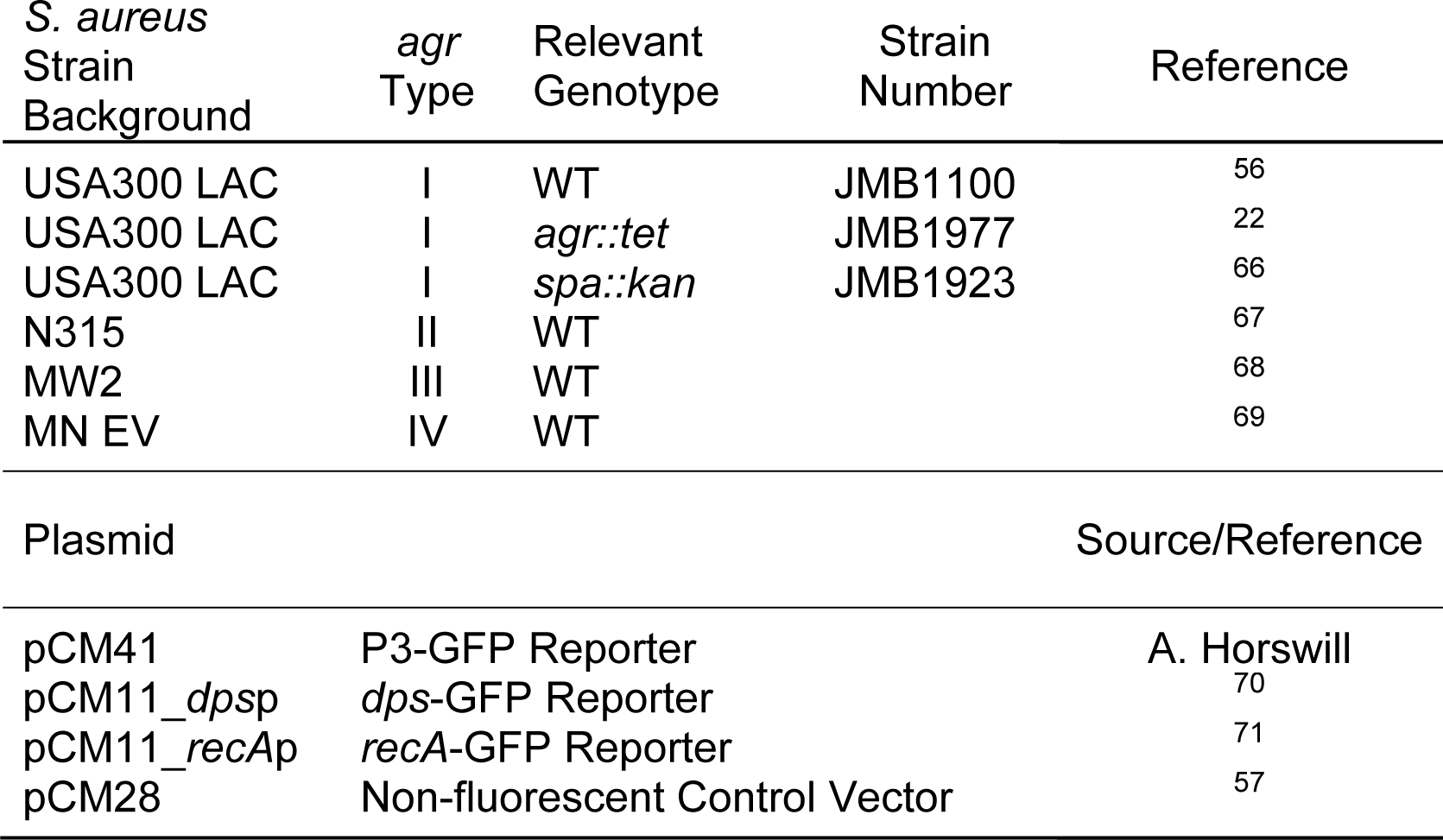
Strains and plasmids

### Growth inhibition assays

Overnight cultures were inoculated into 20-mL tubes containing 5 mL of either TSB, TSB supplemented with 5% v/v DMSO, or 5% v/v DMSO with the indicated concentration of compound to an OD (600 nm) of 0.05. The OD was measured at regular intervals to generate growth curves.

### GFP reporter assays

Overnight cultures were inoculated into 2 mL of TSB supplemented with 0.5% w/v glucose, 5% v/v DMSO, 120 μM compound, and antibiotics appropriate for plasmid retention to an OD (600 nm) of 0.1. Following 8 h of growth, fluorescence and OD were measured using a Thermo VarioSkan plate reader. Green Fluorescent Protein (GFP) fluorescence was measured by excitation at 485 nm and emission at 535 nm.

Data were analyzed by normalizing GFP fluorescence to OD (600nm) and then subtracting the normalized fluorescence of a control strain carrying a non-fluorescent vector (JMB1242, a WT background strain containing the pCM28 vector^57^) from values for experimental cultures to account for background signal. Normalized fluorescence values from three triplicates were averaged and compound-treated samples were compared to those of untreated (DMSO only) samples using a Student’s t-test to determine statistical significance.

### Real-Time Quanitative PCR

Overnight cultures were inoculated into 2 mL of TSB supplemented with 0.5% w/v glucose, 5% v/v DMSO, and 120 μM compound. Following 8 h of growth, aliquots corresponding to 1.0 OD⋅mL were collected. The samples were centrifuged at 16,000 × g for 1 min, pellets were resuspended in RNAprotect Bacteria Reagent (Qiagen, Germany) and incubated at room temperature for 5 min. Cells were pelleted by centrifugation at 16,000 × g for 1 min, RNAprotect Bacteria Reagent was discarded, and samples were stored at −80°C. Cells were re-suspended and washed twice with a lysis buffer consisting of 50 mM Tris, 150 mM NaCl at pH 8.0. To lyse the resupsended cells, lysostaphin was added to 15 µg mL^−1^ and cells were incubated at 37°C for 30 min. RNA was extracted using TRIzol (Invitrogen, CA); contaminating DNA was degraded using a Turbo DNA-free kit (Invitrogen, CA); and cDNA was generated using a High-Capacity cDNA Reverse Transcription Kit (Applied Biosystems, CA). Primers for qPCR were designed using Primer3Plus^58^ and are listed in Table 3. qPCR was performed using a GoTaq qPCR Master Mix kit (Promega, WI) and a QuantStudio 3 Real-Time PCR System (Applied Biosystems, CA). Data were processed and analyzed with REST2009 software^59^.

**Table 3:**
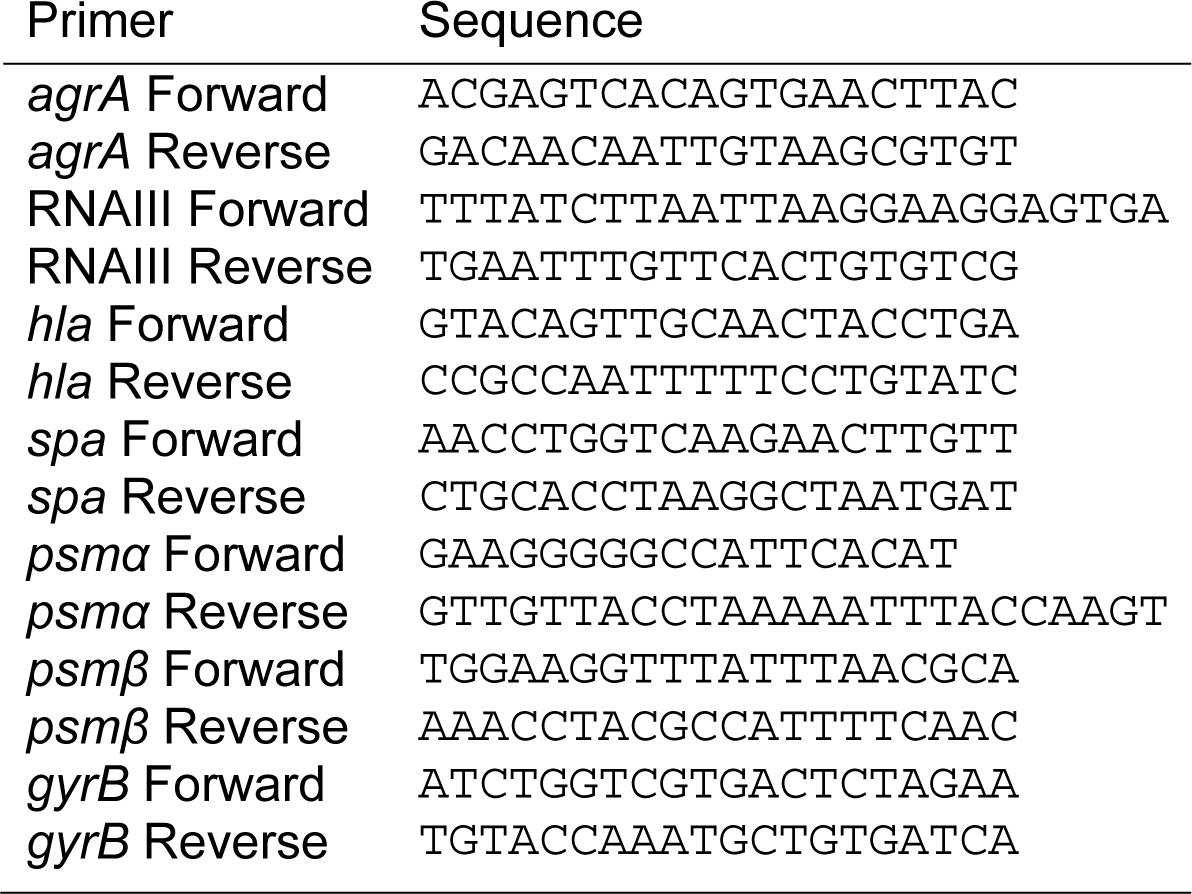
qPCR Primers

### Western blotting

To determine AgrA levels, overnight cultures of USA300 LAC *spa::kan* were inoculated into 25 mL of growth media to an OD of 0.05. Cultures were grown to an OD of ~2.5 at which time aliquots corresponding to 1.3 OD⋅mL were collected. The samples were centrifuged at 16,000 × *g* for 1 min and pellets were washed with 1x PBS before freezing and storage at −20°C. The cultures were then split and compounds were added to a final concentration of 120 μM and 5% v/v DMSO. Samples were again collected after 8 h of growth and processed as described above.

Cell-free extracts were prepared by resuspending cell pellets in 100 μL of a pH 7.6 lysis buffer consisting of 20 mM Tris-HCl, 0.5 mM CaCl_2_, 50 mM NaCl, 40 μg mL^−1^ DNase I, and 20 μg mL^−1^ lysostaphin. The lysis mixture was incubated at 37°C for 1 h, after which 4x SDS-PAGE sample loading buffer spiked with purified AgrA_C_ (for use as a loading control; purification previously described^38^) was added to a final concentration of 1x. Samples were analyzed by electrophoresis using a 15% polyacrylamide SDS-PAGE gel. Protein was transferred to a nitrocellulose membrane, labeled with anti-AgrA_C_ rabbit antisera (Supplementary Fig. S3) and Cy5 goat anti-rabbit IgG secondary antibody (Life Technologies, CA), and visualized using a FluorChem Q Imager. Quantitation of band densities was performed using ImageJ software^60^.

### Isolation of culture supernatants for exoprotein analysis

Culture supernatants for exoprotein analyses (exoprotein profiles, protein quanitation, and hemolytic activity) were obtained simultaneously as described previously^61,62^. Overnight cultures of USA300 LAC and USA300 *agr::tet* were inoculated into tubes containing 5 mL of fresh TSB supplemented with 0.5% w/v glucose, 120 μM compound and 5% v/v DMSO to an OD of 0.1. After 8 h of growth, cell densities were normalized by diluting with fresh media. Cultures were centrifuged at 4,000 × *g* for 20 min to pellet cells and the supernatants were passed through 0.2-μm filters and held at 4°C or frozen at −80°C for long-term storage.

### Exoprotein profiling assay

For SDS-PAGE analysis, supernatants were concentrated by precipitation with trichloroacetic acid by addition to a final concentration of 10% w/v as previously described^61^. Samples were incubated on ice for 30 min before centrifugation at 4,000 × *g* for 20 min. Pellets were washed twice with cold acetone, resuspended in 0.025 the original sample volume of 1x SDS-PAGE loading buffer, and stored at −20°C. Aliquots of triplicates were pooled, analyzed by 15% polyacrylamide SDS-PAGE, and visualized with Coomassie Brilliant Blue. Images were captured using a FluorChem Q Imager.

### Quantitative analysis of secreted protein

Total protein secreted was determined using a Coomassie reagent-staining assay. 5 μL of the filtered supernatant was pipetted into a clear flat-bottom UV plate and 250 μL of 1x Advanced Protein Assay Reagent (Cytoskeleton Inc., CO) was added and mixed. The plate was equilibrated at room temperature for 10 min and absorbances at 595 and 450 nm were measured. The data were processed using the ratio of the absorbance at 595 nm to that at 450 nm for all samples^63^. Ratios for the USA300 LAC *agr::tet* samples were subtracted from the other samples to correct for non-*agr* related exoproteins. The averages of triplicates for samples from cultures treated with compounds were compared to untreated compounds with significance determined by a Student’s t-test.

### Hemolytic activity assay

Hemolytic activity was assessed using a modification of the protocol described by Sully *et al.*^54^. Defibrinated rabbit blood (Hemostat Laboratories) was washed by centrifuging cells at 1000 × *g* for 5 min followed by gently resuspending in ice-cold 1x PBS, and repeating until the supernatant was clear. Blood cells suspended in 1x PBS or dH_2_O were used to determine baselines for no lysis or complete lysis, respectively. The washed red blood cells were added to the plate containing extracts diluted in 1x PBS and controls to achieve final concentrations of 1% rabbit blood and dilutions of culture extract from 1:4 to 1:256. Reactions were mixed by gentle pipetting and incubated at 37°C for 1 h. After incubation, the plates were centrifuged at 1000 × *g* for 5 min at 4°C. 100-µL aliquots of the supernatants were transferred to a 96-well UV-transparent plate and the absorbance at 415 nm was measured. Values of the PBS control were subtracted from the experimental data, and the difference was normalized to the absorbance values from the dH_2_O wells. These values were plotted versus the concentration of extract to generate an activity curve, which was fitted to a four-parameter logistic model using SigmaPlot 10 (Systat Software, Inc., CA). The calculated EC_50_ from the fitting was then inverted to generate the HA_50_. HA_50_ values from cells treated with compounds were compared to those from untreated samples (DMSO only) with statistical significance determined by Student’s t-tests.

### Static model of biofilm formation assay

Biofilm formation was examined as described elsewhere, with minor modifications^61,64,65^. Overnight cultures were diluted into biofilm media^64^ in the presence or absence of compounds, added to the wells of a 96-well plate and incubated statically at 37**°**C for 22 h. Prior to harvesting the biofilms, the OD (590 nm) of the cultures was determined.

The plate was subsequently washed with water, biofilms were heat-fixed at 60**°**C, and the plates and contents were allowed to cool to room temperature. Biofilms were stained with 0.1% w/v crystal violet and destained with 33% v/v acetic acid. The absorbance of the resulting solution was recorded at 570 nm, standardized to an acetic acid blank, and subsequently normalized to the OD of the cells upon harvest.

### AgrA *in vivo* stability assay

Overnight cultures of USA300 LAC *spa*::*kan* were inoculated into 25-mL cultures to an OD of 0.05. The cultures were grown to an OD of 2.0 and samples were collected in volumes equivalent to 1.3 OD⋅mL. Cells from these aliquots were harvested by centrifugation at 18,000 × *g* for 1 min, washed with 1x PBS, and pellets were frozen at −20°C for storage. The remaining cultures were split and erythromycin was added to a final concentration of 10 µM to all cultures except a control. After an incubation of 20 min, samples were again removed and processed. Compounds were added to the remaining cultures to a final concentration of 120 μM and 5% v/v DMSO. Samples were collected after 8 h of growth. All samples were processed for western blotting and analyzed as described above.

### Isothermal titration calorimetry

Purified AgrA_C_ was dialyzed into 20 mM sodium citrate buffer, 250 mM NaCl, and 1 mM (tris(2-carboxyethyl)phosphine) at pH 6.0. The dialysate was retained and used to prepare the compounds. DMSO was added to 5% v/v with an AgrA_C_ concentration of ~375 μM for use with compounds **1** and **2** and 250 μM for use with compound **3**. Compounds **1** and **2** were prepared to concentrations of 10 mM and compound 3 was prepared to 5 mM in the above citrate buffer supplemented with DMSO. Each compound was titrated into the protein solution using a MicroCal iTC 200 Microcalorimeter (Malvern Instruments, UK) at 25°C. The data were processed, analyzed, and fitted using the Origin 7.0 MicroCal module (OriginLab, MA).

## Data availability

All data generated or analysed during this study are included in this published article.

## Acknowledgements

We thank Dr. Alexander Horswill for providing the pCM41 plasmid and the MN EV(406) strain. We also thank Dr. James Millonig and Dr. Paul Matteson for the use of the Real-Time PCR machine and assistance with qPCR experiments. This work was supported in part by the National Institutes Health grant R01GM047958 to AMS, the United States Department of Agriculture MRF project NE-1028 to JMB, and the Charles and Johanna Busch Foundation to JMB. IFB was supported in part by a National Institutes of Health Graduate Training grant (T32 GM008360).

## Author contributions

IFB and AAM performed experiments and analyzed data. IFB wrote the manuscript. All authors designed experiments, reviewed data and reviewed the manuscript.

## Additional information

The authors declare no competing interests.

