## Supplemental Figures S1-S5 for "Drug-like Fragments Inhibit *agr-*Mediated Virulence Expression in *Staphylococcus* aureus"

Figure S1

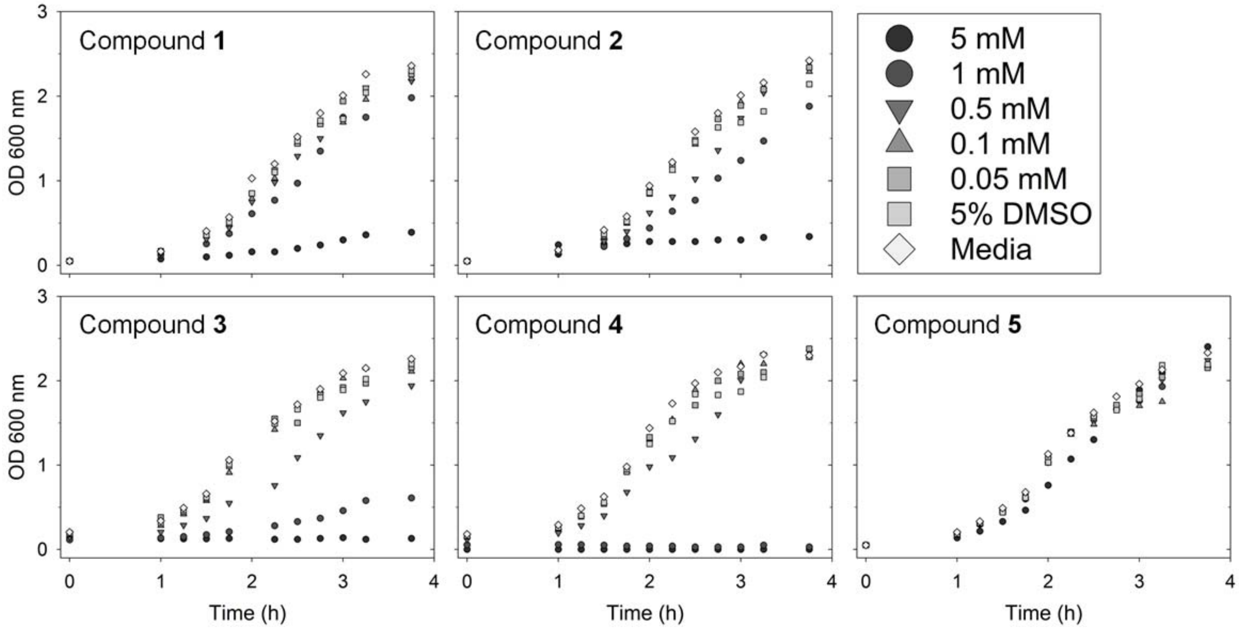

Compound 1
Compound 2
Compound 3
Compound 4
Compound 5
5 mM
1 mM
0.5 mM
0.1 mM
0.05 mM
5% DMSO
Media
OD 600 nm
Time (h)

Figure S1

Compounds do not inhibit growth at concentrations below 0.5 mM. Overnight cultures were inoculated into fresh media containing the indicated concentrations of each compound in 5% DMSO to an OD of 0.1. Cultures were grown with shaking at 37°C, and the OD was measured at regular intervals of time.

Figure S2

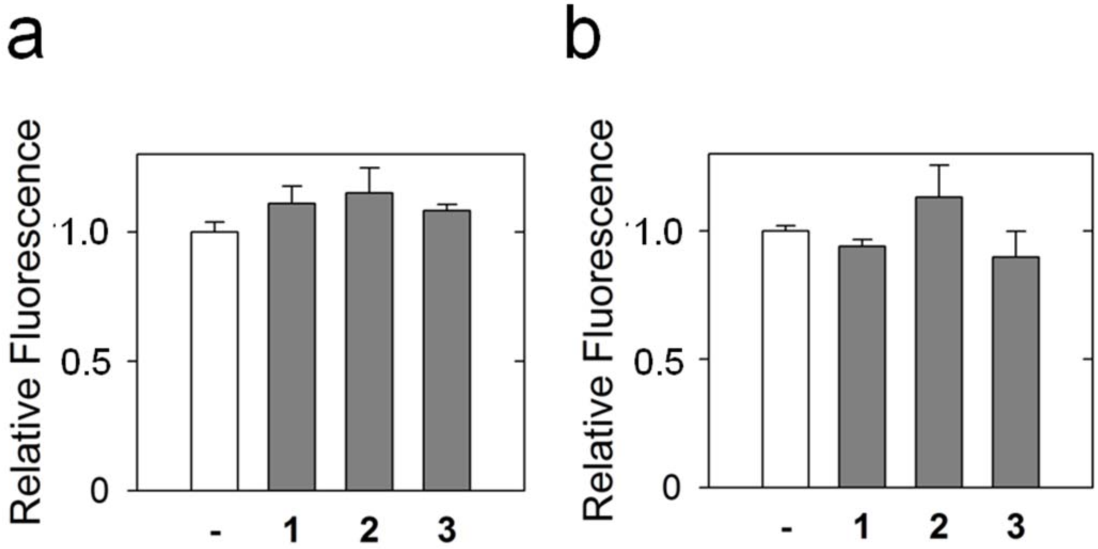
a
b
Relative Fluorescence
1.0
0.5
0
-
1
2
3
Relative Fluorescence
1.0
0.5
0
-
1
2
3

Figure S2

Compounds do not alter transcriptional activity of the *dps* or *recA* promoters. Plasmid-based GFP expression is shown and was driven by *dps* (**a**) or *recA* (**b**) promoters. Cultures were grown for 8 h in the presence of the indicated compound (Cpd.) at a concentration of 120 μM or DMSO alone (**-**). Bars represent OD-normalized fluorescence averaged from three separate experiments relative to the untreated sample (normalized to 1) with error bars indicating standard deviations. Statistical significance relative to the sample without compound was determined using a Student's t-test (* $p<0.05$, ** $p<0.01$).

Figure S3

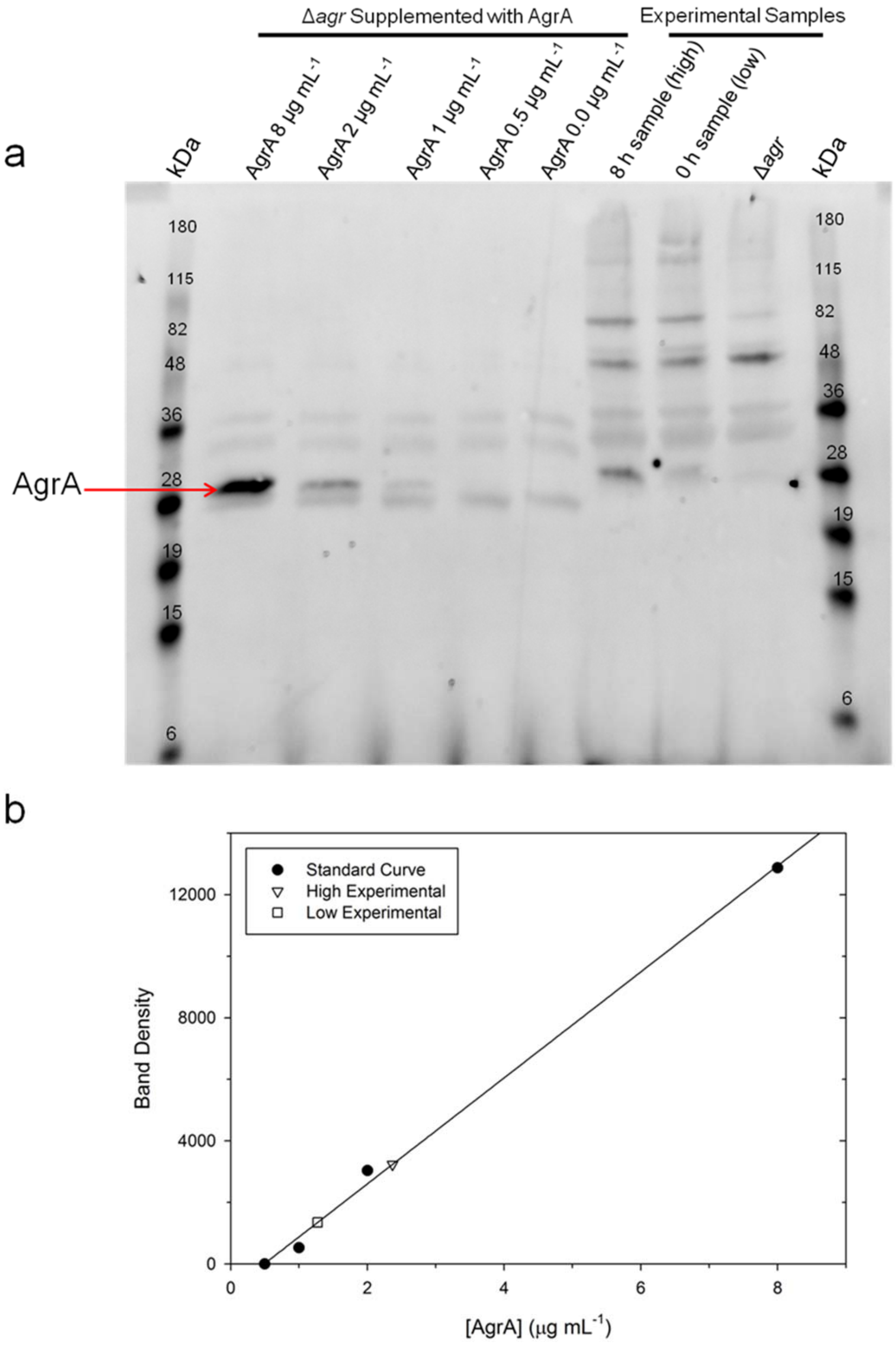

a
Δagr Supplemented with AgrA
Experimental Samples
kDa
AgrA 8 μg mL-1
AgrA 2 μg mL-1
AgrA 1 μg mL-1
AgrA 0.5 μg mL-1
AgrA 0.0 μg mL-1
8 h sample (high)
0 h sample (low)
Δagr
kDa
180
115
82
48
36
28
19
15
6
AgrA
b
Standard Curve
High Experimental
Low Experimental
Band Density
12000
8000
4000
0
0
2
4
6
8
[AgrA] (μg mL-1)

Figure S3

Signal from western blots using anti-AgrA$_C$ rabbit antisera is linear within the range of levels seen in these studies. (**a**) A standard curve for AgrA band intensity relative to protein concentration was generated by adding known quantities of purified AgrA to blot samples generated from Δ*agr* cultures. These samples (15 μl per lane) were analyzed via western blot with along with samples from the experiments reported in Fig. 3a, where a sample taken at 0 h represents the low end of the range of AgrA level from these experiments and the 8 h untreated sample represents the high end. (**b**) Linear regression of the densitometries of the standard curve samples generated the equation y=1722.4x-842.52 with an $R^2$ value of 0.9971. Calculated values from the experimental densitometries fall within the linear range of detection for the tested standards as depicted in (**b**).

Figure S4

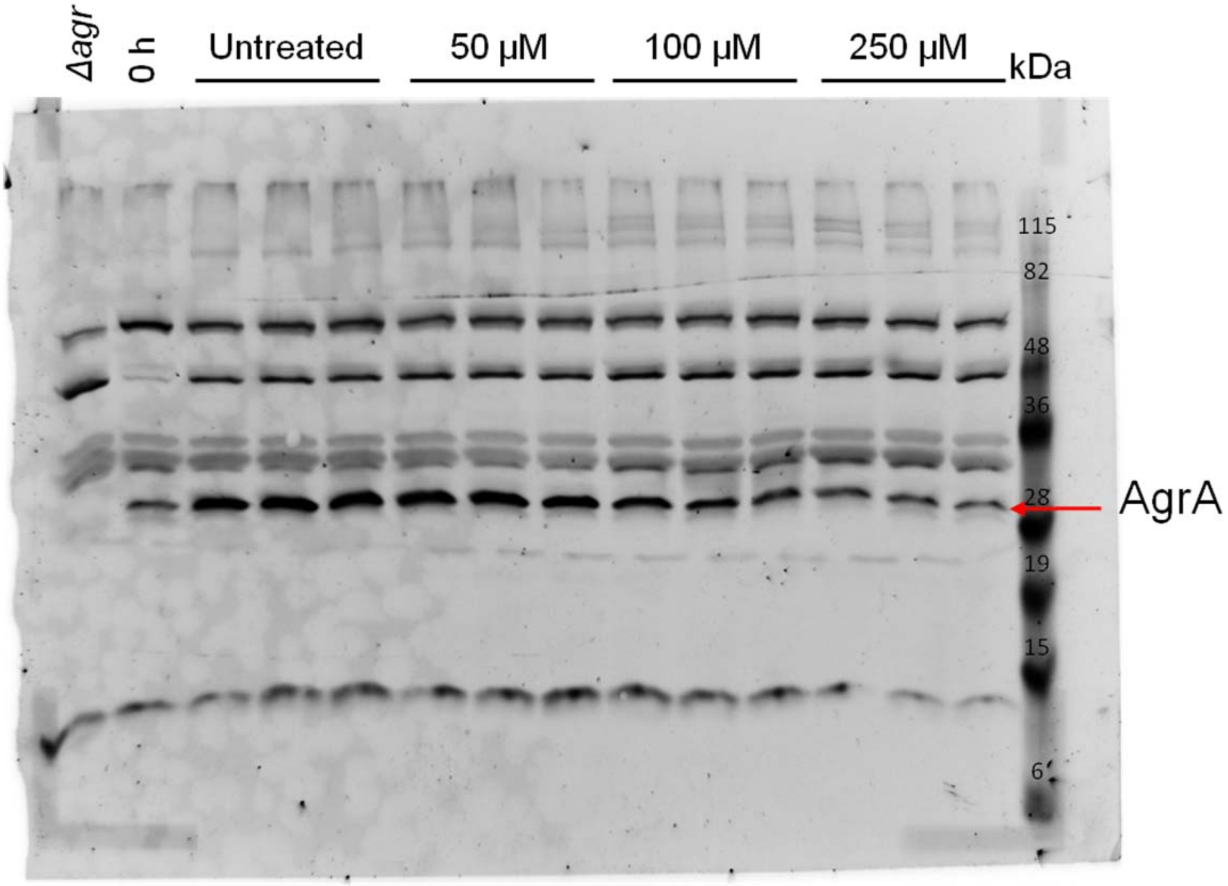
Δagr
0 h
Untreated
50 μM
100 μM
250 μM
kDa
115
82
48
36
28
19
15
6
AgrA

Figure S4

Example experimental western blot.  Cultures of USA300 *spa::kan* were grown to OD 2.5 (0 h) at which time the cultures were divided and compounds were added at the indicated concentrations.  Samples were harvested after 8 h of growth and AgrA levels were determined by western blotting using anti-$AgrA_C$ antisera.

Figure S5

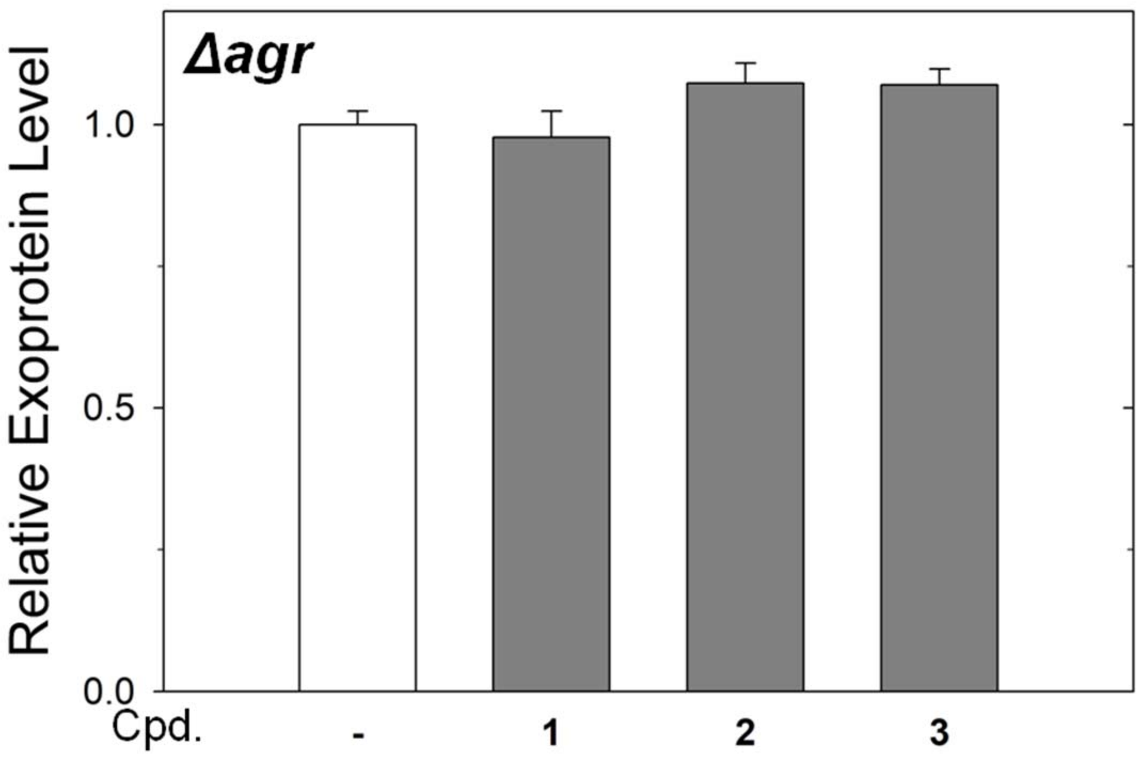

Figure S5

Expression of exoproteins proteins in the Δ*agr* strain is unaltered by compounds. The concentration of total secreted soluble protein was determined using a Bradford protein assay. Levels of secreted protein in the Δ*agr* strain were determined from the average of three replicates. Bars represent average values relative to the untreated sample (normalized to 1) with errors bars representing standard deviations.
